# Dynamics of the p53 response to ionizing and ultraviolet radiation

**DOI:** 10.1101/367995

**Authors:** Elizabeth A. Fedak, Frederick R. Adler, Erin L. Young, Lisa M. Abegglen, Joshua D. Schiffman

**Affiliations:** Department of Mathematics, The University of Utah, 201 Presidents Circle, Salt Lake City, UT 84112; Department of Biology, The University of Utah, 201 Presidents Circle, Salt Lake City, UT 84112; Huntsman Cancer Institute, The University of Utah, 2000 Cir of Hope Dr, Salt Lake City, UT 84103; Department of Pediatrics, Huntsman Cancer Institute, The University of Utah, 2000 Cir of Hope Dr, Salt Lake City, UT 84103; Department of Oncological Sciences, The University of Utah, Salt Lake City, UT 84112

## Abstract

The tumor suppressor protein p53 compiles information about cellular stressors to make decisions on whether the cell should survive or undergo apoptosis. However, the p53 response depends on the source of damage, displaying a ‘digital’ oscillatory response after ionizing radiation (IR) damage and a proportional non-oscillatory response following UV damage. We propose a mathematical model that qualitatively replicates this observed behavior. The difference in p53 dynamics in the model results from two mechanisms: IR damage is fully detected minutes after exposure while UV damage is detected over several hours; and the p53-controlled transcriptional response is dominated by inactive p53 following UV damage. In particular, we hypothesize that an unidentified positive feedback loop controlled by inactive p53 is required to maintain the qualitative high p53 response to UV damage. This work proposes an explanation for two distinct responses of p53 to DNA damage and how each response can lead to cell cycle arrest or apoptosis.

**Author summary:** We propose a mathematical model hypothesizing how the tumor suppressor protein p53 produces two contrasting dynamical responses in response to different types of DNA damage. In particular, we predict the existence of a positive feedback loop controlled by the inactive form of p53, which allows the cell to respond to slowly detected damage. The existence of differing dynamic responses by p53 has implications for our understanding of tumor development and possibly p53-related therapeutic strategies.

## Introduction

Although the functional purpose of p53 within a cell has been widely studied, its dynamics have yet to be fully characterized. The p53 tumor suppressor protein, mutated in 50% of all cancers, is responsible for activating cell cycle arrest or apoptosis programs following cellular stress [1–4]. To guide these decisions, p53 must integrate information about stress from multiple sources—including DNA damage, hypoxia, transcriptional stress, and telomere erosion—that each affect its total level and activation dynamics differently. It is unknown how p53 controls cell cycle arrest and apoptosis through these dynamical changes.

Particularly interesting early work demonstrated that oscillations in total p53 level with a consistent period and amplitude were observed in MCF7 cells exposed to γ–radiation, inspiring a generation of dynamical p53 models [5]. When the same types of cells were exposed to UV light, however, no such oscillations were observed; total p53 instead increased proportional to the amount of induced damage [6]. Both phenomena have also been observed in non-cancerous cells [7, 8]. Researchers have characterized the first response as digital, and the second as proportional [6]. In this work, we use mathematical models to explore what causes this difference in behavior, and further hypothesize why this change in dynamics is necessary for the p53-mediated apoptotic pathway to function after exposure to each type of damage.

Several critical proteins act upstream and downstream of p53 in the apoptotic pathway. We focus on the pathways responsible for detecting double-strand DNA breaks (DSBs) induced by many types of ionizing radiation (IR), and aberrations in DNA structure caused by UV radiation, collectively known as UV photoproducts [9]. DSBs are detected by an aggregate of Mre11, Rad50 and Nbs1 (known as the MRN complex) minutes after damage [10, 11]. Of the UV photoproducts, about 85% are cyclobutane pyrimidine dimers (CPDs), and about 15% are 6–4 photoproducts (6–4PPs) [12, 13]. 6–4PPs and CPDs on the transcribed strand of DNA are both detected and repaired quickly, while CPDs on the non-transcribed strand remain unrepaired after several hours [14–16]. Undetected photoproducts can cause additional DSBs or single-strand DNA breaks (SSBs) when the DNA transcription machinery attempts to process a damaged strand [12].

Both types of damage are detected by cellular pathways that communicate with p53 through phosphorylation of ataxia telangectasia mutated (ATM) and ataxia telangectasia mutated related (ATR) proteins [15]. A 2004 study suggests UV-mediated activation of ATR only happens after replication stress, but does not address ATM, and a more recent study shows ATM being upregulated 4–8 hours after UV exposure by DSB formation [17, 18]. These two kinases can phosphorylate p53 on Ser15 (a state which we call p53-arrester), modifying its transcriptional activity; and phosphorylate Mdm2, the primary regulator of p53, on Ser394, targeting it for autoubiquitylation [6, 19]. This downregulation of Mdm2 allows cellular p53 levels to rise, promoting transcription of p53 binding targets. We give special attention to two transcriptional targets: PTEN, which sets off a cascade sequestering Mdm2 in the cytoplasm through suppression of Akt; and Wip1, a phosphatase which acts to deactivate members of the p53 apoptotic pathway once damage is repaired [20–22]. Wip1 has been shown to dephosphorylate both ATM_*p*_ and ATR_*p*_ [23].

Mathematical models of p53 damage response have considered the IR and UV damage responses of p53 separately, using the fact that p53 does not oscillate in response to UV damage to simplify dynamics, or accounted for the difference by assuming Wip1 and ATR_*p*_ do not interact [6, 24]. Previous work has centered around p53-Mdm2 oscillation, attributing it to Mdm2 overregulation, Wip1-ATM_*p*_ downregulation, spatial dynamics or stochasticity [25–29]. This paper instead asks how and why p53 levels rise proportionally to UV damage level while oscillating after IR exposure.

By creating a system which can replicate the p53 response to both types of damage, we develop several new predictions about how the p53 pathway works. We first propose that the difference in downstream damage response to UV and γ–radiation damage arises because the p53 pathway responds quickly to IR damage, but slowly to UV damage. Furthermore, in order to properly respond to UV damage, we predict the existence of a stabilizing feedback loop released by inactive p53 and suppressed by active p53. These predictions suggest the oscillatory and non-oscillatory responses may aid the cell in responding to both slowly and quickly detected DNA damage.

## Methods

### Model

The full model builds on the model posed in Zhang *et al*. [24]. To disregard cell cycle arrest and apoptosis, we have removed the model components related to phosphorylation of p53 on Ser46 (MAP3K, MAP3k_*p*_, MAP2K, MAP2K_*p*_, MAP2K_*_p_p*_, p38MAPK, p38MAPK_*p*_, p38MAPK_*pp*_, HIPK2, p53AIP1, p53DINP1) and indicators of cell cycle arrest or apoptosis (p21, CytoC, Casp3). We add new components to track DNA damage products and their detection mechanisms (DSB, CPD_*t*_, CPD_*td*_, CPD_*nt*_, CPD_*ntd*_). Due to the scaled nature of the fitting data, all quantities are nondimensional; we choose parameters based on quantitative results when available, and focus on implications of the quantitative dynamics.

Suppose the number of UV photoproducts produced is proportional to the magnitude of UV exposure, scaling the damage level such that [CPD_0_] = 10 represents exposure to 10 J/m^2^ UV. We let [CPD_*t*_] represent the number of unrepaired CPDs on the transcribed strand, [CPD_*nt*_] represent the number of unrepaired CPDs on the non-transcribed strand, [CPD_*td*_] and [CPD_*ntd*_] represent transcribed strand and non-transcribed strand CPDs in the process of being repaired, respectively; [DNAP] represent cellular levels of DNA polymerase, [DSB] represent the number of DSBs in the cell, and [ATM_*p*_] represent the concentration of active ATM. We assume damage is distributed evenly between transcribed and non-transcribed strands, such that 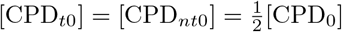.

CPDs on the transcribed strand are detected (*k_ctd_*) and repaired (*k_ctr_*) quickly. CPDs on the non-transcribed strand are detected slowly (δ*k_ctd_, δ*< 1), and can become DSBs if they are not repaired before the DNA strand is split by replication machinery [16]. The cell is assumed to be in S phase for the duration of the simulation. Here, we assume the speed of nucleotide excision repair (NER) and DNA replication is limited by the amount of available DNA polymerase in the cell [38]. DNA polymerase can repair CPDs on either type of strand (CPD_*td*_ or CPD_*ntd*_), be free, or be stalled at a double-strand break during replication ([DSB]). We normalize the total available amount of DNA polymerase to 1, such that 
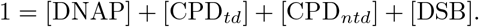

The activation of p53 due to ATR_*p*_ is assumed to be negligible, with p53 activation dominated by ATM_*p*_/ATR_*p*_ response to replication-induced DSBs. Earlier versions of the model accounted for the detection of both CPDs and DSBs, with ATM_*p*_ upregulating p53 when double-strand breaks were present and ATR_*p*_ upregulating p53 if CPDs were present. This led to unresolvable discrepancies between CPD detection and repair rates and the initial delay in post-UV p53 levels observed in Batchelor et al., 2011 (Figures S2 and S3) [6]. We do not track the amount of MRN complex in the cell, as DSB detection happens on a very fast timescale. Furthermore, despite the evidence that p53 is able to upregulate the rate of DNA repair, we assume flagged sites of DNA damage are repaired at a constant rate. Under these assumptions, the damage detection module is as follows: 
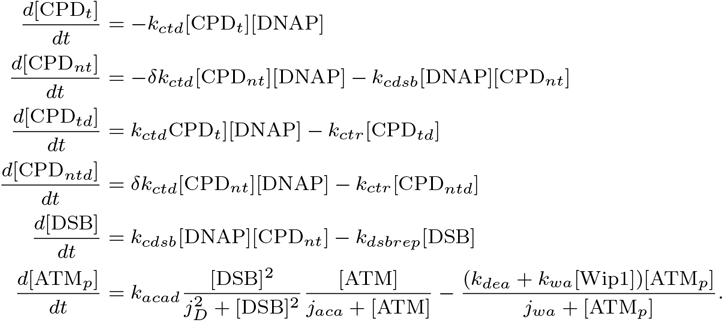

The second-order Hill function term for DSB upregulation of ATM mimics spatial dynamics of rate-limiting and scarcity. When the number of DSBs is high, the cell can only detect them at a maximal rate limited by the amount of MRN complex; and when the number of DSBs is low, the remaining DSBs have a lower probability of being detected at a given time. ATM_*p*_ deactivation is modeled as a Michaelis-Menten term because it is carried out by phosphatases such as Wip1.

Downstream of ATM_*p*_, we consider p53 and its transcriptional targets. p53 exists in the model in two classes: unmodified (p53) and Ser15-phosphorylated (p53_*a*_). Nucleic p53 upregulates transcriptional targets by binding in tetrameric form to their promoter regions [39]. Since p53 dimerizes cotranslationally, concentrations of p53 are understood to be concentrations of p53 dimers, where tetramers are dimers of dimers [39]. Binding to promoter regions therefore occurs at a rate proportional to [p53]^2^, [p53_*a*_]^2^, or both. Tetramers of p53 and p53_*a*_ are ignored in this work. In accordance with experimentally observed p53 behavior, unmodified p53 is produced at a high, constant rate, then quickly ubiquitylated by nucleic Mdm2, after which it is degraded or shuttled to the cytosol. Mdm2, the primary regulator of p53, is split into three classes: cytosolic Mdm2 (Mdm2_*c*_), nucleic Mdm2 (Mdm2_*n*_), and Ser394-phosphorylated nucleic Mdm2 (Mdm2_*np*_). Mdm2 is assumed to be quickly shuttled to the cytosol once produced. To model p53 upregulation of Mdm2 without tracking Mdm2 mRNA, we let *P* be the probability that p53 is bound to the promoter region of Mdm2, *k_on_*_1_ be the rate of binding of a single inactive p53 dimer to the Mdm2 promoter region, *k_on_*_2_ the rate of binding of a single active p53 dimer to the same region, and *k_off_* be the dissociation rate of tetramers from the promoter region. Then 
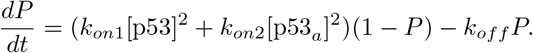

If the probability equilibrates quickly, we estimate 
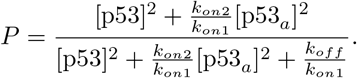

This term is scaled in the full model by combined transcription/translation rate *k_tm_*, with 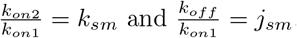.

Once sequestered in the cytosol, Mdm2 cannot affect nucleic p53 levels until phosphorylated on Ser186 by Akt_*p*_, promoting its transfer to the nucleus. Nucleic Mdm2 is responsible for downregulating nucleic p53, but can also be phosphorylated on Ser394 by ATM_*p*_, targeting it for autoubiquitylation and subsequent degradation.

Phosphorylated Mdm2 can be rescued from its fate by spontaneous dephosphorylation or by Wip1 interference.

The other transcriptional targets of p53, PTEN and Wip1, are upregulated by p53 in a rate-limited manner proportional to p53^2^ or p53^2^_*a*_, respectively. Two other components of the Akt/PTEN pathway are included in the system: PIP_3_, which can be formed from PIP_2_ at a substrate-limited rate and dephosphorylated by PTEN, and Akt_*p*_, which is activated by PIP_3_ [40]. Wip1 is capable of dephosphorylating ATM_*p*_, Mdm2_*np*_, and p53_*a*_, helping return the system to a pre-damage state.

We assume no change in total ATM, PIP2, or Akt concentration, and nondimensionalize the total available amount of each protein to 1, such that the conservation equations 
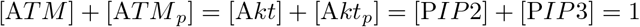
account for the active and inactive states of each protein. We also assume PTEN equilibrates quickly once its transcriptional switch is flipped by an increase in p53 concentration, and normalize by the ratio of transcription rate to decay rate, such that PTEN is represented in the full model by 
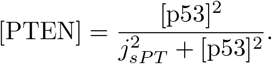

The three conservation equations, along with this system of differential equations, form the transcriptional response module: 
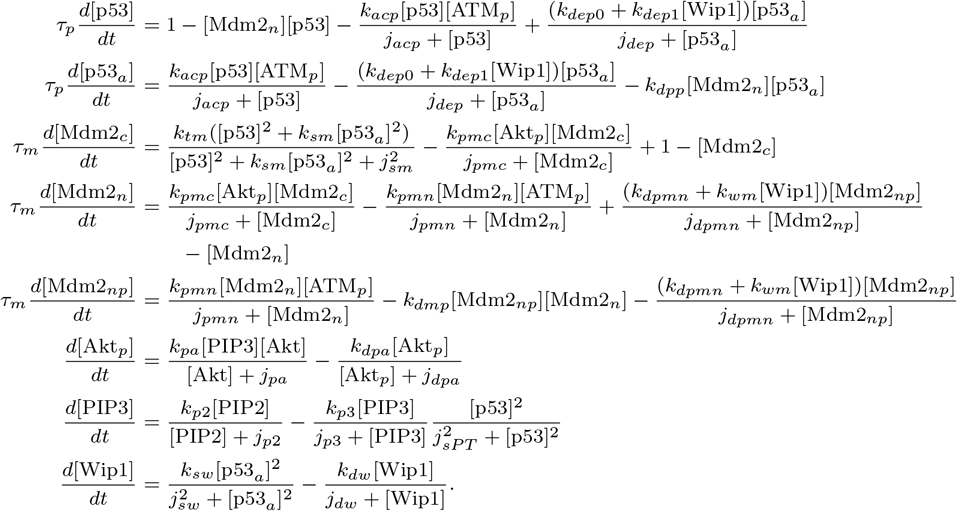

Michaelis-Menten forms are used for enzymatic reactions, where all terms are rate-limited by substrate. Parameter values, units, sensitivity analysis, and descriptions can be found in Table 1 of the Supporting Information, along with details of the nondimensionalization.

### Numerical methods

Numerical solutions to these systems of ODEs were obtained using Matlab’s built-in fourth-order Runge-Kutta algorithm, `ode45`. Bifurcation diagrams were obtained with XPPAUT, and plotted in Matlab using the `plotbd` files written by Maurizio De Pitta and available through the MathWorks website. Fits to experimental data were optimized using an MCMC simulated annealing algorithm. For the CPD fitting, we minimized the least-squares distance of the damage response module from Fig. 6B in [30]. To fit total p53 response to UV damage, we minimized the sum of least squares distances from the model with [CPD_0_] set to 2, 4, 6, 8 or 10 to all 5 respective trials in Fig. 1f of [6]. For the IR response, we minimized the sum of the least squares distances from the model period, amplitude, and mean to the respective average period, amplitude and mean of Fig. 1C in [6].

## Results

### Damage detection module

All proteins included in the model can be found in Figure 1. The damage detection module is highlighted in green.

**Figure 1.**
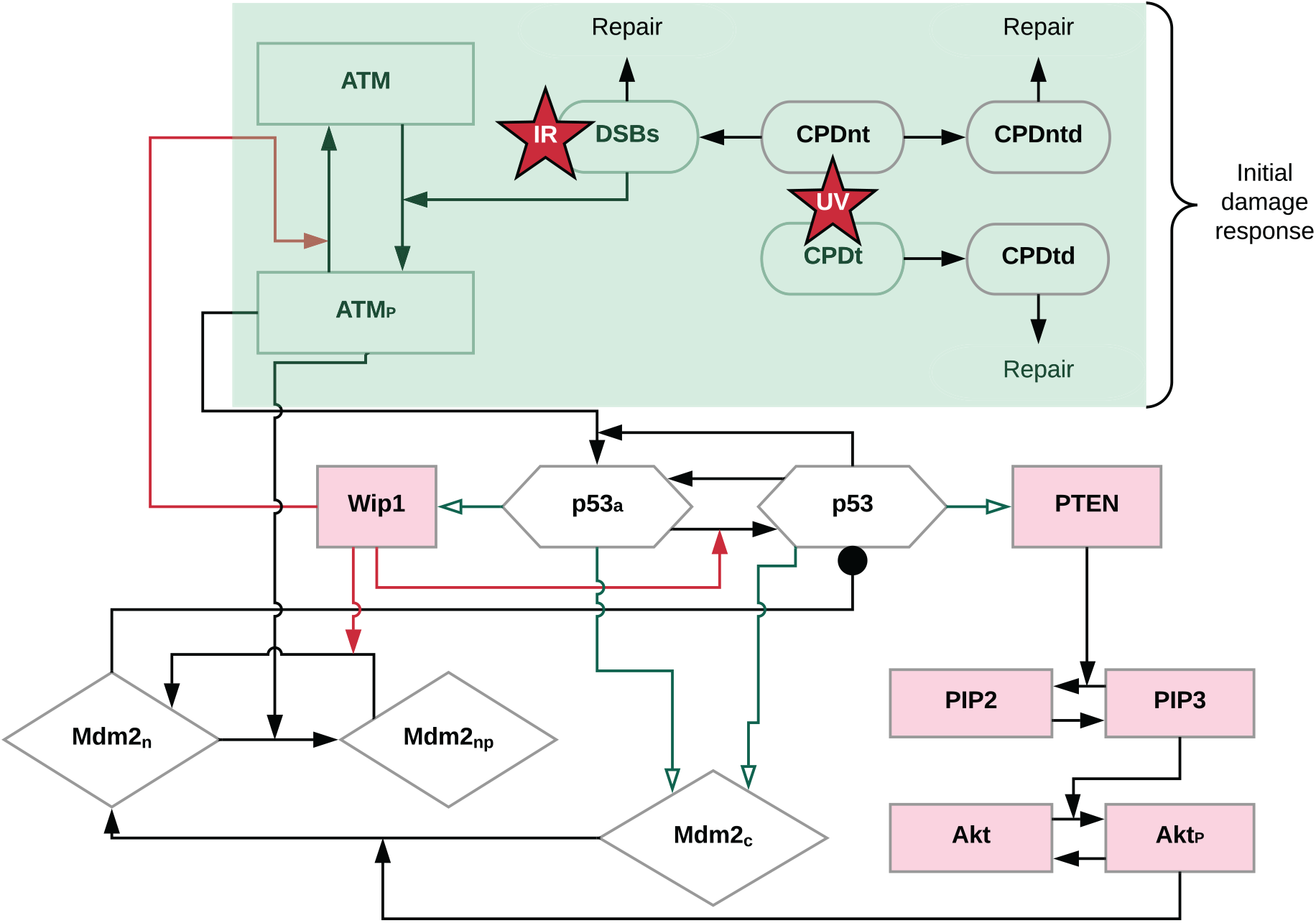
Diagram of the full model. Green arrows represent transcriptional upregulation by p53, red arrows represent dephosphorylation by Wip1, and circles represent inhibition. During the initial damage response (light green), we assume γ–irradiation produces double-strand breaks (DSBs), while UV damage distributes CPDs between the transcribed and non-transcribed strands (CPD_*t*_ or CPD_*nt*_). These either are repaired, or, if the cell is in S phase, non-transcribed strand CPDs have a chance of becoming DSBs and upregulating the p53 pathway. The remainder of the diagram shows the transcriptional response module. Inactive p53 self-regulates by promoting transcription of Mdm2, but can also activate the PTEN pathway. Active p53 promotes transcription of both Mdm2 and Wip1. Mdm2 is quickly shuttled to the cytosol once translated, and remains there until it is phosphorylated on Ser186 by active Akt. Nucleic Mdm2 downregulates p53 and is susceptible to downregulation by active ATM. Wip1 is transcriptionally upregulated by active p53, and can dephosphorylate active p53, active ATM, and Ser394-phosphorylated Mdm2 tagged for autoubiquitylation. Proteins highlighted in pink have been left out of the simplified model.

Fibroblast data suggest that CPDs are initially repaired quickly, but that after the percentage of CPDs remaining in the cell after 8 hours is much higher than we would predict from an exponential model with a constant repair rate [30]. Furthermore, in MCF7 cells, the post-UV p53 response is delayed by 30 minutes to 5 hours [6]. Both sets of data suggest that not all UV damage is treated equally: some portion of damage is detected and repaired quickly without upregulating the p53 pathway, and some damage evades initial detection and repair but upregulates the p53 pathway. If we assume quickly detected CPDs can upregulate p53 through ATR_*p*_, it is impossible to induce the observed delay in p53 induction. Conversely, if we assume all CPDs are detected slowly, p53 upregulation can be delayed—but only in response to a delay in CPD detection, which conflicts with experimental observations. Hence we develop a damage detection model which considers at least two classes of CPDs: transcribed strand CPDs, which are detected and repaired quickly; and non-transcribed strand CPDs, which are detected and repaired slowly [16, 31]. Assuming ATR_*p*_ and ATM_*p*_ equally upregulate the p53 pathway removes the benefit of this construction, as the ATR-dependent mechanism still receives the largest signal before the observed peak of p53 activity.

We resolve the above issues by assuming that photoproducts induced by UV damage are distributed evenly between transcribed and non-transcribed strands, and that ATM interaction with DSBs dominates ATR interaction with CPDs in p53 activation. We thus expect p53 to be upregulated only when DSBs are created from unrepaired CPDs during DNA replication when the cell is in S phase. Variability in the delay in p53 upregulation in MCF7 cells can therefore be explained by considering that cells may be in different stages of the cell cycle [6].

By separately handling transcribed strand ([CPD_*t*_] and [CPD_*td*_]) and non-transcribed strand CPDs ([CPD_*nt*_] and [CPD_*ntd*_]), where undetected non-transcribed strand CPDs have a chance of becoming DSBs during DNA replication, we were able to capture both the experimentally observed CPD repair dynamics (Figure 1 of SI) and the delay in p53 upregulation. This construction allows for a delay in ATM_*p*_ induction after UV damage, but for an immediate upregulation of ATM_*p*_ following *γ*–irradiation damage, simulating the difference in signal exposure speeds. Predicted ATM_*p*_ levels are treated as input to one of the two p53 pathway models outlined below.

### Simplified model

We first explore whether it is possible to model both types of damage response without involving transcriptional targets of p53. This simplified model ignores Akt/PTEN and Wip1 feedback loops, but does allow Mdm2 to be upregulated by p53. In this model, Mdm2 suppression of p53 was the only negative feedback loop mechanism capable of producing oscillations after IR damage, and the change in ATM_*p*_/ATR_*p*_ induction was sufficient to remove oscillations after UV damage (Figure 2). However, if Mdm2 overactivity provides the oscillating mechanism, oscillations will occur whenever the p53 concentration passes a certain threshold. Oscillations after UV damage can therefore only be avoided by ensuring p53 stays below this threshold. Producing a non-oscillatory UV response led to a tenfold difference in average p53 concentration between the IR (Figure 2A) and UV (Figure 2B) cases. This is inconsistent with data showing that total post-UV p53 can be sustained at higher levels and for longer durations than the average IR spike [6, 7]. Because this is impossible to achieve if Mdm2 overactivity causes oscillations whenever p53 crosses a threshold, we hypothesize that additional p53 targets must be involved in creating the two distinct dynamical responses.

**Figure 2.**
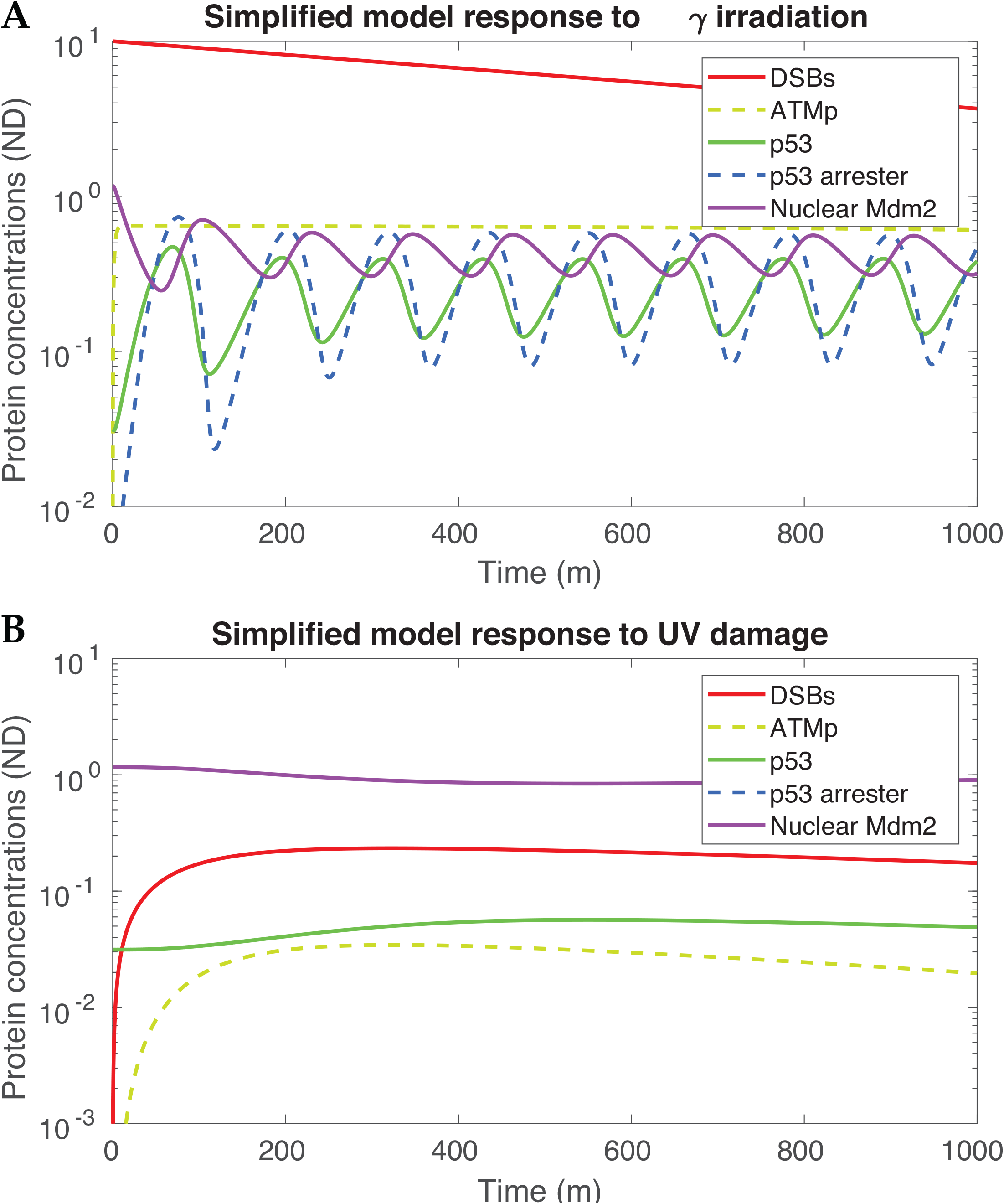
A demonstration of the failure of the simple model, which ignores the downstream transcriptional targets of the p53 pathway, to properly capture post-UV dynamical response. Oscillations in this system are caused by Mdm2 overactivity (A), but this same mechanism over-suppresses total p53 concentration when the system is exposed to a slowly detected signal (**B**). Here, γ irradiation is modeled by setting [DSB]_0_ = 10, UV radiation by setting [CPD]_0_ = 10.

### Transcriptional response module

The transcriptional response module incorporates the positive feedback loop caused by p53 upregulation of PTEN and the negative feedback loops caused by p53 upregulation of Wip1. PTEN sets off a chain of events that sequester newly transcribed Mdm2 in the cytosol [32, 33]. This creates a bottleneck for replenishment of nucleic Mdm2, and thus slows ubiquitination of nucleic p53 even as Mdm2 is translated at higher rates. Wip1 is a phosphatase which targets active ATM ([ATM_*p*_]), active p53 ([p53_*a*_]), and nucleic phosphorylated Mdm2 ([Mdm2_*np*_]) [22, 34].

To differentiate the transcriptional responses to slowly and quickly detected damage, we use ATM_*p*_ as a bifurcation parameter rather than as a state variable. For very low concentrations of ATM_*p*_, we do not expect the p53 pathway to be significantly upregulated. However, if ATM_*p*_ levels rise due to a slowly detected source of damage, we expect ATM_*p*_ to pass some threshold at which the p53 response is turned on. If the source of damage is quickly detected, we expect ATM_*p*_ to rise to high levels almost instantaneously, after which it may interact with Wip1 in the full model. Therefore, we expect a bifurcation diagram of change in p53 equilibrium concentrations with respect to ATM_*p*_ to have three distinct regions: one showing p53 inactivity when ATM_*p*_ is low, one showing the fast-damage transcriptional response when ATM_*p*_ is high, and one showing the slow-damage transcriptional response in between.

Mediating the bifurcation profile of the transcriptional module (Figure 3) allows the creation of a regime in which p53 releases to high levels in response to UV damage, but oscillates in response to IR (Figure 4). Here, the equilibria can be split into three classes: low, which is stabilized either by a lack of damage signal or the regulatory processes of Wip1; high, stabilized by sufficiently high levels of PTEN; or an intermediate saddle connecting the high and low equilibria.

**Figure 3.**
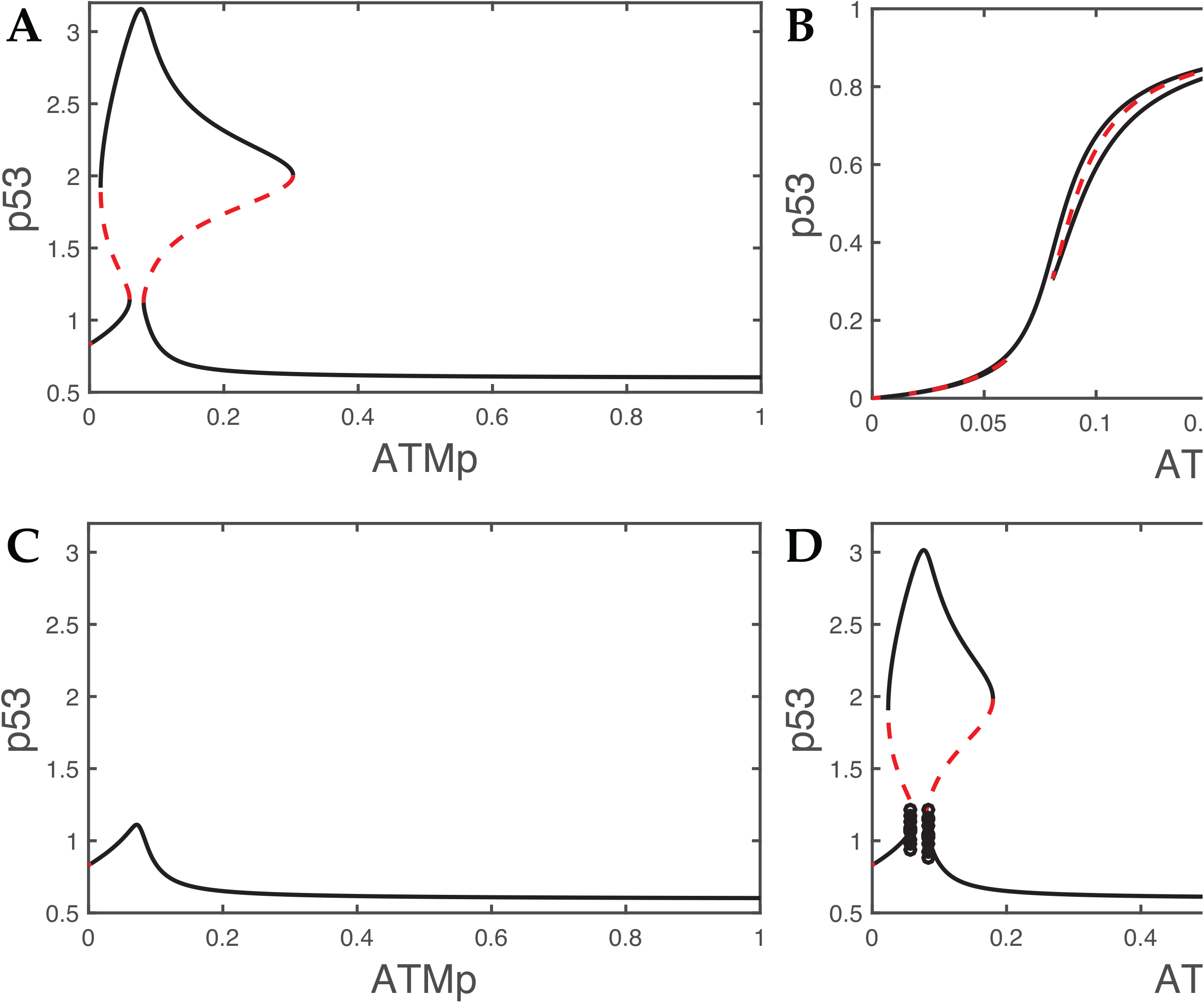
Bifurcation diagrams demonstrating the intermediate ATM_*p*_ release mechanism for p53 (A) and p53_*a*_ (B). Uses best-fit parameters to [6], *k_pmc_* = 32 and *k_dm_* = 0. 1 (where *k_pmc_* is the rate of nucleic import of Mdm2 and *k_dm_* the basal decay rate of Mdm2). This structure takes on topologically diverse forms depending on the values of the parameters: in **C**, we set *k_pmc_* = 34, connecting the two low equilibrium branches and collapsing the high equilibrium; and in **D**, we set *k_dm_* = 0. 0009. A Hopf bifurcation occurs on the low equilibrium branch, decoupling the low equilibrium destabilization from the limit points.

A system with no induced DNA damage stays at low equilibrium levels of p53. The low p53 equilibrium collapses at medium levels of ATM_*p*_ by a novel codimension-2 bifurcation, such that the p53 concentration tends towards high equilibrium in this intermediate region (Figure 3A). When ATM is activated slowly, as in the case of UV exposure, p53 stays on the low branch until [ATM_*p*_] = 0.6, upon which it is drawn towards the high branch until all damage is repaired. We expect p53_*a*_ concentrations to be roughly proportional to the concentration of active ATM_*p*_; however, once the system stabilizes to high p53 equilibrium, increased levels of substrate (p53) compensate for lower levels of activating enzyme (ATM_*p*_). Active p53 is therefore present in higher levels on the high branch than it would be if there were no window of escape (Figure 3B).

However, when damage is detected quickly, ATM_*p*_ rises to a region where the low equilibrium is stable before p53 levels can escape to high equilibrium. Furthermore, high concentrations of ATM_*p*_ effectively convert p53 to p53_*a*_, which in turn releases Wip1. In the full system, Wip1 would then interact with ATM_*p*_ to induce oscillations on p53 level corresponding to a back-and-forth motion on the right branch of the low equilibrium. This mechanism is notably robust when the damage detection module is changed. Since the only requirement on the input is the experimentally observed conclusion that IR-induced damage is detected quickly and UV-induced damage is detected slowly, we observe the same differential dynamic profile in models accounting separately for ATM_*p*_ and ATR_*p*_ (Figure S4).

We make additional assumptions about active p53 behavior to guarantee existence of this behavioral regime. For the PTEN pathway to stabilize at medium levels of phosphorylated ATM, and for the unstable window of escape to exist, PTEN must be preferentially transcribed by inactive (unphosphorylated) p53. Likewise, Ser15-phosphorylated p53 must be more proficient at upregulating transcription of the negative regulatory components of the system, Mdm2 and Wip1.

### Full model

The full model contains both the damage response module and the transcriptional response module. By exposing a system with the above transcriptional bifurcation structure to damage, we were able to induce conditions under which p53 increases to high levels in response to UV damage, but oscillates in response to IR (Figure 4). The same equations and parameters were used to create both diagrams, with the exception of DSB repair rate (*kdsbrep*). Since DSBs in the UV damage case are created by stalled repair machinery, they are expected to be repaired significantly faster (*kdsbrep* = 0. 0087) than DSBs created by *γ* irradiation (*kdsbrep* = 0. 004).

**Table 1.** A list of key assumptions and predictions made in this work.

| Key observations | Key assumptions | Key predictions |
| --- | --- | --- |
| • The cell repairs CPDs at two different rates, and p53 activation is delayed following UV damage<br>• p53 responds qualitatively differently to IR and UV damage<br>• Total p53 levels post-UV damage can be higher than total p53 levels post-IR | • There are multiple mechanisms for UV damage handling<br>• IR and UV damage are detected at different rates by the p53 pathway<br>• By upregulating different proteins, inactive p53 can self-stabilize, but active p53 self-regulates | • DSBs activate the p53 pathway much more strongly than CPDs<br>• Having two dynamic regimes prevents the cell from overreacting to quickly detected damage, or from underreacting to slowly detected damage<br>• A positive feedback loop stabilizes inactive p53 |

**Figure 4.**
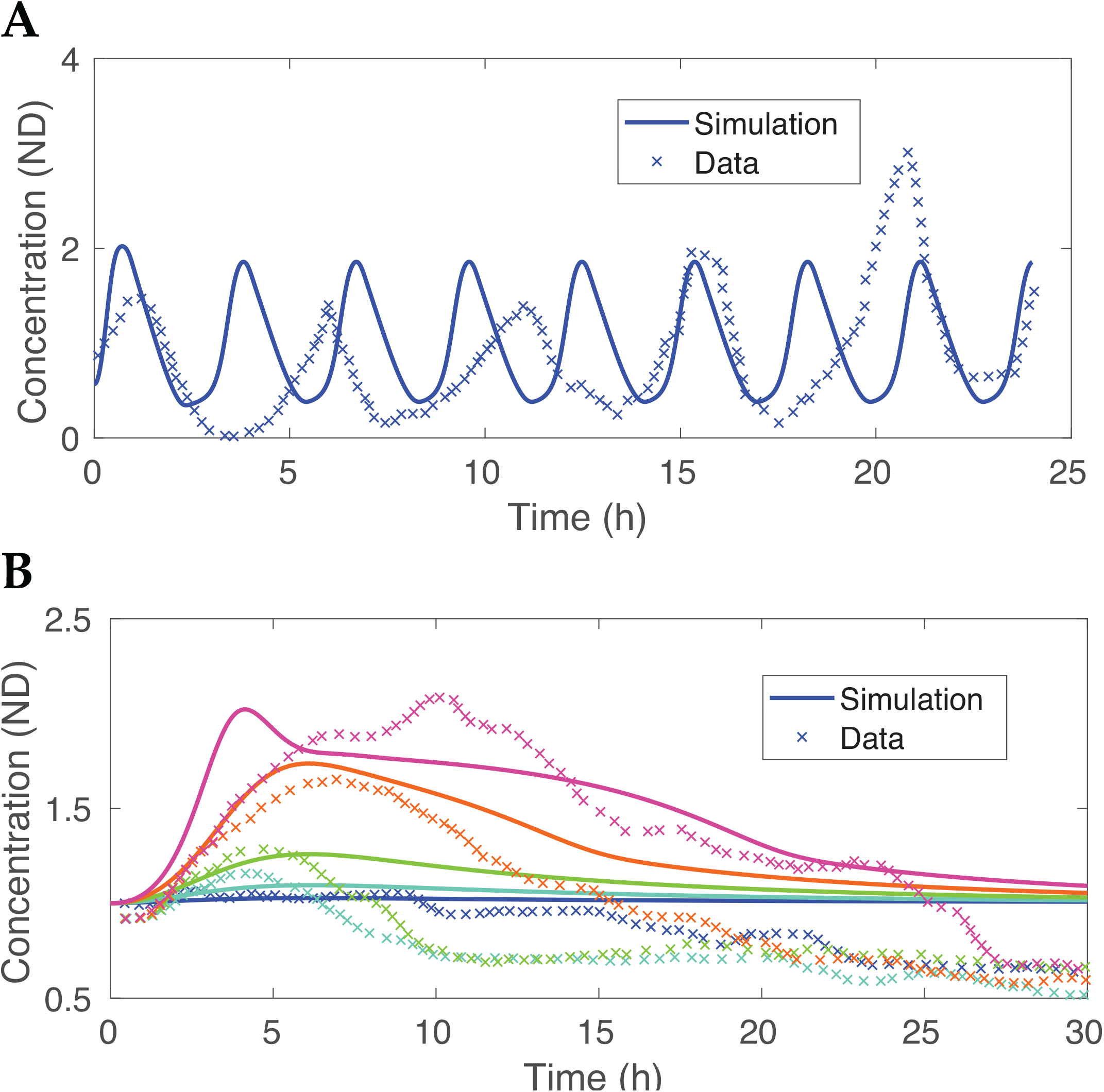
Model predictions compared to data from [6]. **A** Response to γ-irradiation normalized to equilibrium, [DSB]_0_ = 5. Of the three illustrative cases provided, the average period was 3.548 hours and the average amplitude was 2.395; here, the model converges to a solution with a period of 2.97 hours and amplitude 1.424. Changing the initial DSB concentration by a factor of 2 did not impact the period or magnitude of oscillations. **B** Normalized response to UV radiation, demonstrating the proportional post-UV p53 response (combined LS error = 28.75). DSB damage repair (*kdsbrep*) is enhanced, as DSBs created during this process stall replication machinery and thus localization for repair would be faster.

## Discussion

### Mechanistic implications

This model introduces a robust mechanism that causes p53 to oscillate in response to ionizing radiation and respond proportionally to UV damage.

The difference in behavior is caused, in part, by a difference in upstream signal strength. Damage caused by ionizing radiation is detected within minutes: this leads to the stabilization of active p53, which upregulates Wip1, which in turn deactivates ATM_*p*_, producing oscillations. In contrast, UV photoproducts are detected and repaired over a period of hours, leading to activation of a positive feedback loop that stabilizes inactive p53. The oscillations can be understood as the p53 regulatory network responding to overstimulation, then overregulation, in a series of pulses. Slowly detected damage, however, is unable to stimulate protein activation quickly enough to trigger oscillations. p53, as a damage aggregator, should be capable of making decisions on cell fate regardless of the rate at which damage is detected. If the p53 response is quickly upregulated in response to ionizing radiation, a cell runs the risk of overactivating the p53 pathway in response to a manageable number of DSBs. Conversely, if the cell is exposed to a slowly detected source of damage, the reduced concentrations of p53-activating kinases could cause a failure to respond to even crippling levels of damage. We propose that the differential dynamical responses of p53 arise to address this variation in detection speed. A cell exposed to small amounts of *γ*–irradiation would activate ATM_*p*_ and thus active p53 in high concentrations, but if no signal is detected after Wip1 is turned on, the damage response stops. Instead, the regulatory pathways are only activated if the system spikes multiple times, as has been suggested in earlier works [35]. The oscillations in total p53 level may then be seen as a mechanism for avoiding overreactions to repairable levels of DSBs. In contrast, the cellular response to UV photoproducts depends more on the duration than the magnitude of the signal: if the cell sustains medium levels of ATM_*p*_, the amount of active p53 produced will depend on the amount of time the cell spends on the high branch of equilibrium. Hence the stabilization mechanism for intermediate signaling levels ensure the cell can respond to slowly detected sources of damage.

For this intermediate-signal release to occur, we predict the existence of a self-stabilizing feedback loop triggered by inactive p53. That is, to compensate for the reduced rate of p53 activation caused by a deficit of ATM_*p*_, the cell increases concentrations of inactive p53 through transcriptional regulation. No such mechanism is needed if ATM_*p*_ persists in high concentrations. Hence, we predict two distinct mechanisms of p53 release capable of regulating the cell cycle: one characterized by low levels of inactive p53 and high levels of kinase, and one by high levels of inactive p53 and low levels of kinase. It should also be noted that inactive p53 is capable of acting as a transcription factor, but less is known about its function than active p53 [36].

The necessity of a slow signaling response, the presence of oscillations, and the ability of a cell to recover after DNA damage is repaired support the paradigm that active p53 is self-regulatory while inactive p53 is self-stimulatory. Not only does active p53 preferentially transcribe regulatory genes such as *PPM1D* and *MDM2*, a more subtle form of regulation occurs in which removing inactive p53 also removes its stabilizing effect.

### Future work

There is nothing to currently suggest that the Akt/PTEN pathway is indeed the positive feedback loop releasing p53 at intermediate levels of ATM_*p*_. Some earlier work suggests PTEN is preferentially upregulated by Ser46-phosphorylated p53, but not Ser15-phosphorylated p53, and suggests nothing about inactive p53 [21]. To address this problem, we plan to conduct experiments on the binding of p53 to the promoter regions of *PTEN, Wip1* and *Mdm2*, and also consider IGF-BP3, which can perform the same role PTEN has in this model [21].

The bifurcation structure displayed in the transcriptional module is too rich to explore fully in this work. We have successfully derived a simpler model with the same behavior and characterized the unusual codimension 2 and codimension 3 bifurcations.

Lastly, the data suggest several improvements which could be made to fully capture the dynamics of p53. Stochastic effects could be incorporated to resolve the changes in p53 oscillation period and amplitude that cannot be explained by this model, and further regulatory mechanisms of p53 must be considered to explain the drop in p53 level to below equilibrium post-UV damage [6]. These mechanisms may involve tracking the replication phase of the cell, as the cell in our model is assumed to always be in S phase. We also observe that the MCF7 cells used in experiments in [6] do not necessarily converge to the high equilibrium; this is consistent with the observation that MCF7 cells are deficient in apoptotic response, but measuring the p53 response to UV damage in non-pathological cell lines would help to further elucidate the structure of the transcriptional response in non-cancerous cells. To account for variance in p53 oscillation period across species, one could vary time-dependent parameters, notably the nondimensional characteristic speed of p53 activity, *τ_p_* [37]. The model cannot account for the variance in amplitude shown in single cells, nor are the regulatory actions of Mdm2 and Wip1 sufficient to produce the post-damage dip in p53 concentration shown in Fig. 1f of [6]. Continued efforts by our group and others on this model of the dynamic response of p53 activation may lead to a better understanding of p53 function, leading to diagnostic and perhaps one day clinical applications for patients with cancer.

## Acknowledgments

We thank the CHPC at the University of Utah and the Huntsman Cancer Institute for providing computing resources, James Keener for insight on bifurcation analysis, and Chee Han Tan for E.A. Fedak’s personal Matlab license.

## Supporting Information

**S1 Fig. Results of fitting the model to Fig. 6B, [30]** Using MCMC, we estimate *δ* = 0. 042, *k_ctd_* = 0. 0055 and *k_ctr_* = 0. 145 (least squares distance 0.0300).

**S2 Fig. Best fit of simple model CPD detection and repair rates to Fig. 6A of [30], control.** The delay caused by incorporating the positive feedback loop with XPC creates an irreconcilable delay in the repair rate of CPDs.

**S3 Fig. Best fit of CPD detection and repair to model (??) to Fig. 6A of [30], control; and initial fit of total p53 level using model (??) as the CPD detecton and repair module.** Since ATR could still be activated by transcribed strand CPDs, it was not possible to reconcile the fast CPD detection and repair rates with the delay in p53 induction shown in [30].

**S4 Fig. Images from E.A. Fedak, *Dynamics of the p53 response to ultraviolet and ionizing radiation*, Poster session presented at SMB Annual Meeting, 2017, Salt Lake City, UT. A** In response to normalized ionizing ratdiation damage, total p53 levels oscillate using the earlier ATR_*p*_-dependent model. B In response to different levels of UV damage, total p53 level rises proportionally to the amount of damage caused.

**S5 Fig. Parameter sensitivity analysis demonstrating change in goodness-of-fit to all levels of UV damage in Batchelor 2011.** All parameters listed have been increased or decreased by 10% to measure the percent change in least squares fit.

**S6 Fig. Parameter sensitivity analysis demonstrating change in goodness-of-fit to the average period, amplitude, and mean of oscillations in Fig. 1c of Batchelor 2011.** All parameters listed have been increased or decreased by 10% to measure the percent change in least squares fit.

**S7 Fig. Parameter sensitivity analysis demonstrating change in goodness-of-fit to only the average period in Batchelor 2011.** All parameters listed have been increased or decreased by 10% to measure the percent change in least squares fit.

S1 Table. Parameters used in the simple model.

S2 Table. Parameters used in the CPD detection module.

S3 Table. Parameters used in the nondimensional full model.

S1 File. An expanded description of models and methods.

